# A physiological Na^+^ signal activates mitochondrial PINK1 to couple synaptic depression to caspase-3-dependent spine pruning

**DOI:** 10.64898/2026.05.29.728915

**Authors:** Benjamin Chun-Kit Tong, Inmaculada Segura, Simone Wanderoy, Markus Rehm, Ariane Metzger, Melanie Boerries, Angelika Bettina Harbauer

## Abstract

Mitochondrial density in dendrites adapts synaptic inputs to sustain local ATP and Ca^2+^ buffering. During long-term depression (LTD), synapse elimination is accompanied by activation of caspase-3 through sublethal mitochondria-derived apoptotic signals. However, upstream signals that link synaptic activity to mitochondrial remodeling remain unknown. Here we show that Na^+^ influx through NMDA receptors depolarizes mitochondria during chemically induced LTD. This triggers stabilization and activation of PINK1 in a translation-dependent manner, leading to asynchronous mitochondrial fission. Na^+^ influx and PINK1 are required for cLTD-induced fission, and blocking either prevents caspase-3 activation and spine shrinking in cultured neurons. Together, these findings identify a Na^+^-PINK1 signaling that couples NMDA receptor activity to mitochondrial fission and caspase-3-dependent synapse elimination during LTD, with implications for the homeostatic regulation of synaptic density.

## Introduction

Synaptic plasticity is a fundamental cellular mechanism that enables neurons to reshape their connectivity by either growing new connections or by removing others. Two opposite paradigms termed long-term potentiation (LTP) and long-term depression (LTD) have been particularly well studied, which increase or decrease the number of synaptic inputs, respectively (*1*, *2*). Mitochondria have been shown to adapt to the increased synaptic demand during LTP, by polarizing and moving towards the postsynaptic sites (*3*) as well as by altering their ultrastructure and association with the cytoskeleton (*4–6*). Directly after LTP induction, a synchronized burst of mitochondrial fission occurs in the postsynaptic site due to CamKII-dependent phosphorylation and activation of the mitochondrial fission GTPase DRP1 (*6*). Blocking of mitochondrial fission prevents dendritic spine growth upon LTP induction, revealing the importance of mitochondrial support for synaptic growth. Fittingly, the proportional volume of mitochondria in dendrites within the hippocampal formation *in vivo* mirrors the density of synaptic input, which depends on regulation of mitochondrial fission by Ca^2+^-induced signaling (*7*).

During LTD, mitochondria play an equally important role. As gatekeepers of the pro-apoptotic molecule cytochrome C, mitochondria control the non-apoptotic activation of the protease caspase-3 (*8*, *9*). Cytochrome c release occurs through mitochondrial outer membrane permeabilization (MOMP), which can be mediated by the pore-forming BCL-2 family proteins BAX, BAK, and BOK (*10*). Among these proteins, BAX has been shown to be required for NMDA receptor-dependent LTD and the associated non-apoptotic activation of caspase-3 (*11*). Consistently, Caspase-3 deficient organisms lack structural remodeling of neurons (*8*, *12–14*). Intriguingly, this non-apoptotic role of caspase-3 for LTD is abolished in PTEN-induced kinase 1 (PINK1) knockout mice (*15*). As a central coordinator of mitostasis (*16*), PINK1 is perfectly positioned to initiate mitochondrial downscaling in response to LTD, yet a mechanistic connection between the induction of LTD and PINK1 activation is still missing. PINK1 activation in distal neurites requires its local translation (*17*) as well as its stabilization at the translocase of the outer membrane (TOM complex, (*18*)) to prevent its rapid degradation after cleavage by PARL (*19–21*). Experimentally, PINK1-dependent mitophagy can be induced by the depolarization of mitochondria with ionophores like CCCP or inhibitors of the respiratory chain that generates the electrochemical gradient (*19*, *20*), yet a physiological stimulus of PINK1 activation has not been identified.

Here we show that induction of LTD in cultured hippocampal neurons leads to a transient depolarization of mitochondria due to Na^+^-influx through the NMDA receptor. This leads to the stabilization and activation of PINK1 at the outer mitochondrial membrane, driving caspase-3 activation and mediating spine shrinkage. This provides a physiological stimulus for PINK1 activation in the absence of mitochondrial damage.

## Results

### cLTD induces asynchronous dendritic mitochondrial fission distinct from cLTP

To visualize synaptic changes and mitochondrial remodeling during synaptic plasticity, we performed time-lapse microscopy of primary hippocampal neurons transfected with cytosolic GFP and mitochondria-targeted mScarlet. Consistent with previous reports (*4*, *22*), dendritic mitochondria in mature neurons at DIV14 appeared as long, tubular structures with limited mobility along the entire dendritic shaft (Supplementary Fig. 1a,b). Neurons were stimulated either with glycine or N-methyl-D-aspartate (NMDA) to induce chemical long-term potentiation (cLTP) or chemical long-term depression (cLTD), respectively, verified by dendritic spine morphological changes (Fig. 1a). Notably, mitochondrial fission was observed in both cLTP and cLTD (Fig. 1a,b, white arrowheads, and supplementary movie 1), despite the opposing effects of these plasticity paradigms on dendritic spine morphology (Fig. 1a,b, yellow arrowheads). Simultaneous application of the NMDA receptor antagonist D-AP5 abolished fission, confirming that mitochondrial fission in both paradigms was driven by NMDA receptor-dependent ion influx (Fig. 1a,b). Induction of cLTD led to a small but significant decrease in mitochondrial occupancy along dendrites (Fig. 1c), fitting to a downscaling of mitochondria in anticipation of a reduction in synapse numbers.

**Fig. 1.**
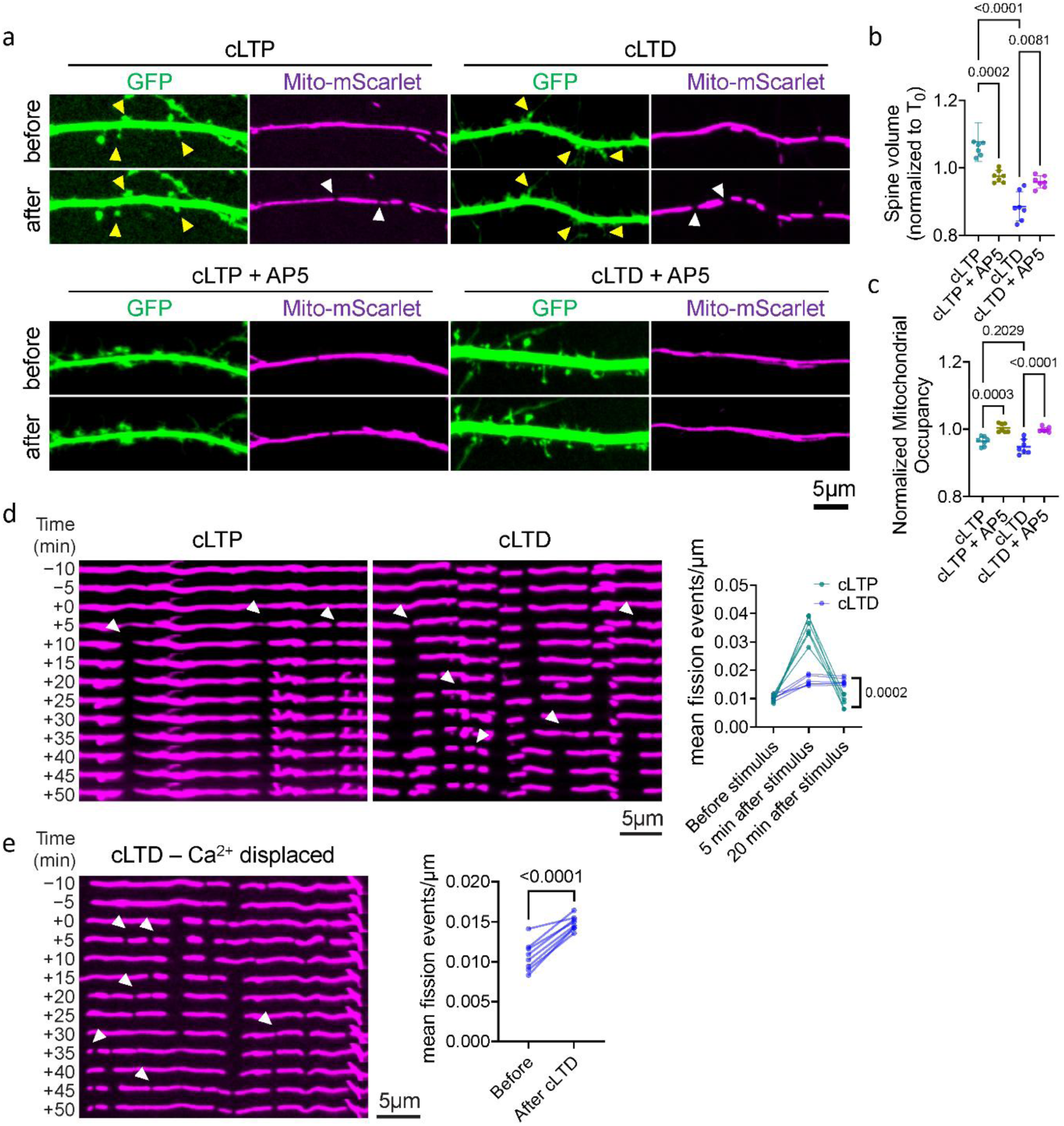
cLTD induces sustained dendritic mitochondrial fission in a manner distinct from cLTP. **a**, Representative images of dendritic spines and mitochondria in hippocampal neurons before and after induction of chemically induced long-term potentiation (cLTP; 400 µM glycine) or long-term depression (cLTD; 20 µM NMDA), with or without co-application of the NMDA receptor antagonist D-AP5 (AP5, 40 µM). Yellow arrowheads indicate dendritic spines undergoing morphological changes. White arrowheads indicate mitochondrial fission events. See also supplementary movie 1. **b**, Quantification of spine volume before and after cLTP or cLTD stimulation, with or without D-AP5 (40 µM). Spine volume was quantified from 7 experiments. Each point represents one neuron. Statistical analysis was performed using one-way ANOVA followed by Tukey post hoc tests. **c**, Quantification of mitochondrial occupancy in dendrites before and after cLTP or cLTD stimulation. Mitochondrial occupancy was quantified from 7 experiments. Each point represents one neuron. Statistical analysis was performed using one-way ANOVA followed by Tukey post hoc tests. **d**, Representative kymographs of dendritic mitochondria during cLTP and cLTD. Summary plots show mitochondrial fission events over time following stimulation. Mitochondrial fission events were quantified from 6 experiments. Each point represents one neuron. Statistical analysis was performed using unpaired Student’s t-test at 20 min after stimulation. White arrowheads indicate mitochondrial fission events. **e**, Representative kymographs and quantification of mitochondrial dynamics during cLTD under Ca^2+^-free conditions. Mitochondrial fission events were quantified from 8 experiments. Each point represents one neuron. Statistical analysis was performed using paired Student’s t-test. White arrowheads indicate mitochondrial fission events.

Intriguingly, while cLTP triggered a rapid, transient burst of mitochondrial fission, cLTD induced asynchronous mitochondrial fission over an extended period of time (Fig. 1d). A sustained elevation in fission rate could be observed even 20 min after stimulation (Fig. 1d). This suggested that the mechanisms underlying mitochondrial remodeling during cLTP and cLTD were not identical. As fission during cLTP depends on Ca^2+^ influx, we tested whether extracellular Ca^2+^ was required for mitochondrial fission in both plasticity paradigms. Unexpectedly, mitochondrial fission during cLTD persisted under Ca^2+^-free conditions (Fig. 1e), confirming that it is mechanistically and temporally distinct from cLTP-associated mitochondrial fission.

**Supplementary Fig. 1.**
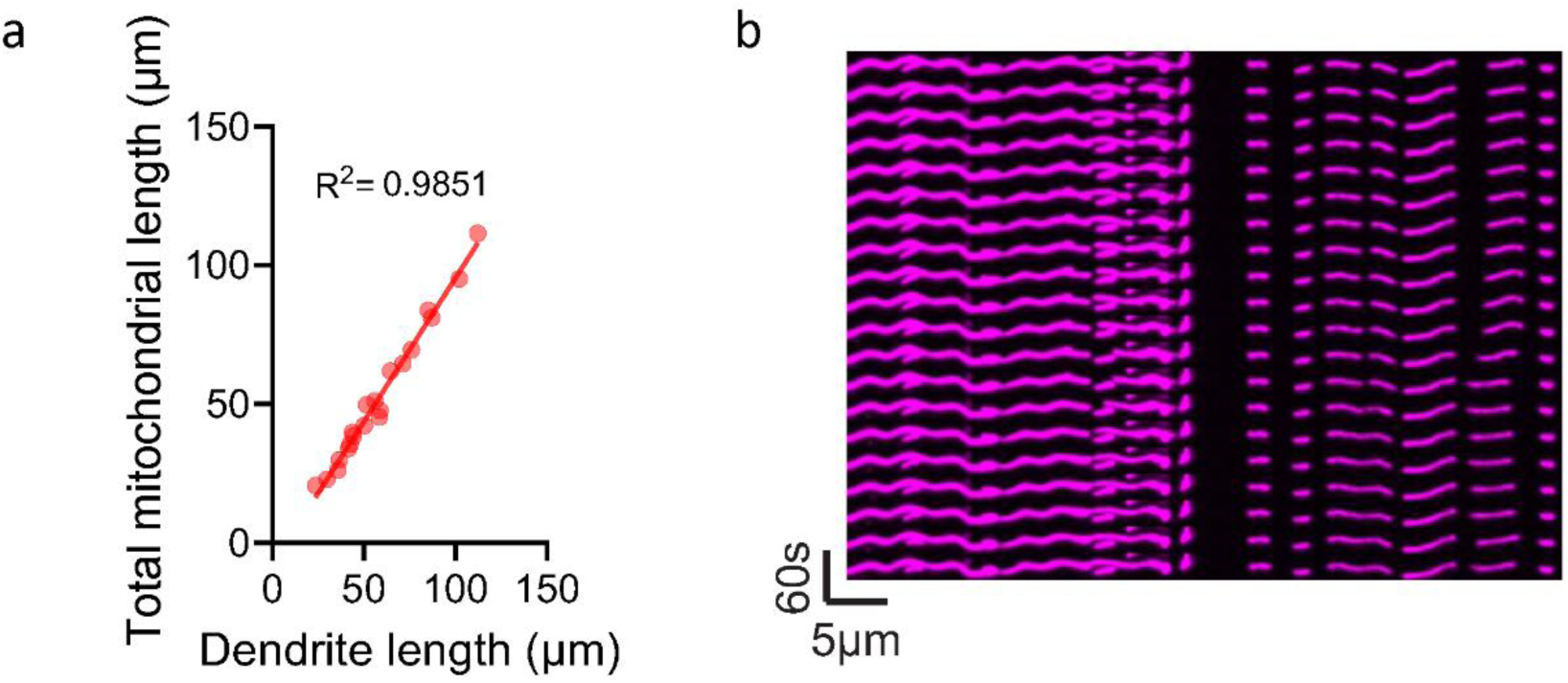
Dendritic mitochondria are spatially stable. **a**, Scatter plot of total mitochondrial length versus dendrite length. Each point represents one dendritic segment. **b**, Representative time-lapse kymograph of mito-mScarlet–labeled mitochondria in a dendrite of a DIV14 mouse primary hippocampal neuron under control conditions.

### cLTD induces reversible Na+-dependent mitochondrial depolarization

To understand whether mitochondrial remodeling observed during cLTD was associated with changes in mitochondrial function, we targeted the FRET-based biosensor ATeam (*23*) to the mitochondrial matrix to assess mitochondrial ATP levels (Supplementary Fig. 2a,b). Fluorescence lifetime-based imaging of the changes in donor lifetime (FLIM-FRET (*24*), validated by application of the complex III inhibitor Antimycin A, Supplementary Fig. 2c), revealed that acute treatment with NMDA led to a transient decrease in mitochondrial ATP levels (Fig. 2a). This change was quickly compensated, indicating that NMDA triggers a reversible change in mitochondrial bioenergetics. This was accompanied by a rapid drop in mitochondrial membrane potential (measured with the potential sensitive dye TMRE, Fig. 2b). Mitochondrial depolarization was followed by gradual recovery, suggesting that it does not reflect irreversible mitochondrial damage or toxicity (Fig. 2b). Mitochondrial depolarization could be blocked by D-AP5, confirming that it was driven by NMDA receptor activation rather than nonspecific effects of NMDA application (Fig. 2c).

**Fig. 2.**
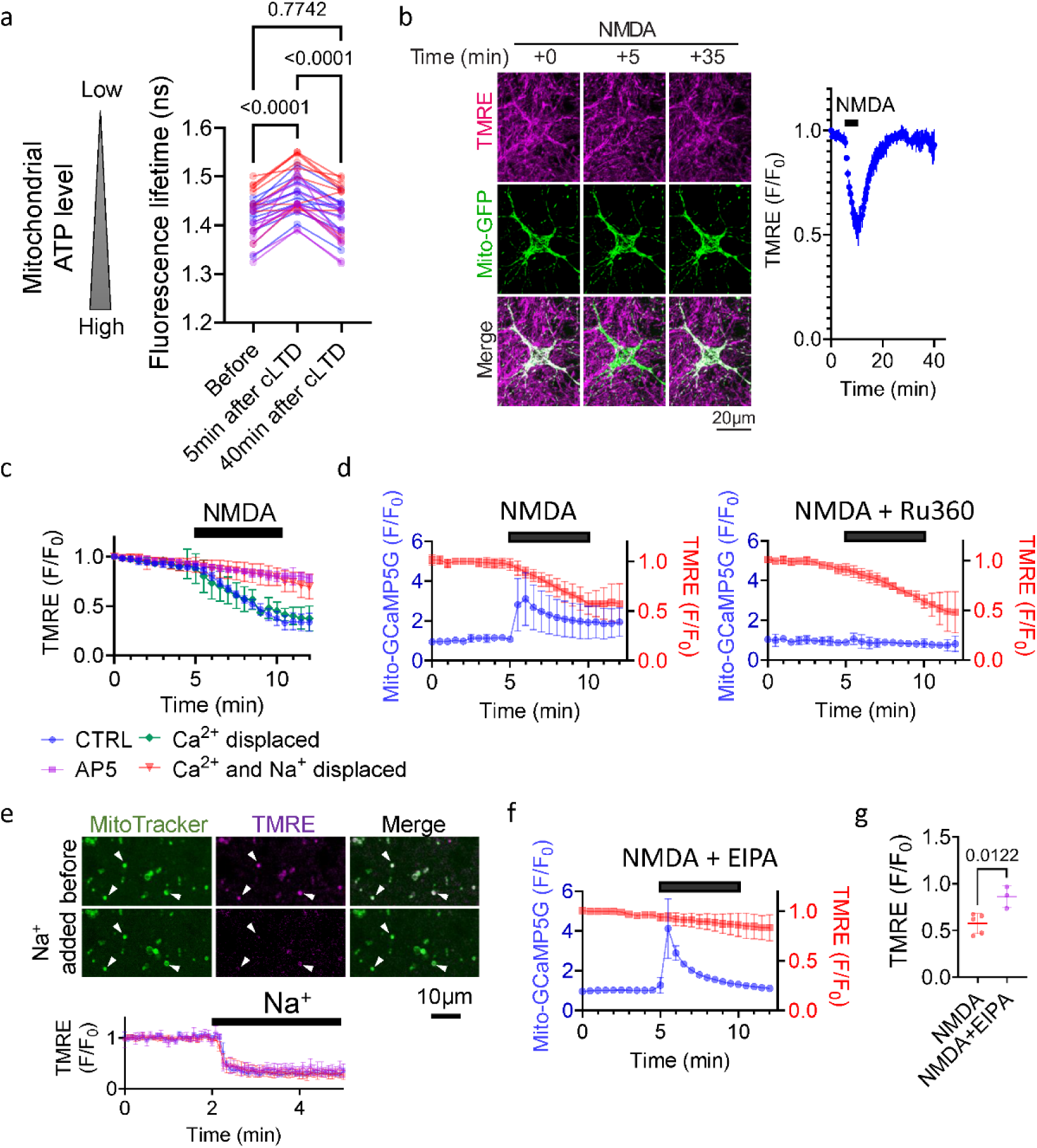
NMDA induces rapid but reversible, Na^+^-dependent mitochondrial depolarization. **a**, Evaluation of mitochondrial ATP levels using the mitochondrial-localized FRET-based biosensor mito-ATeam before and after cLTD induction. Mitochondrial ATP levels were quantified from 31 neurons from 3 experiments. Each point represents one neuron. Statistical analysis was performed using repeated-measures one-way ANOVA followed by Tukey post hoc tests. **b**, Representative images and quantification of mitochondrial membrane potential using TMRE in hippocampal neurons following cLTD induction. Data were quantified from 4 experiments. **c**, TMRE-based quantification of mitochondrial membrane potential under different ionic and pharmacological conditions during cLTD. Conditions include extracellular Ca^2+^-free medium, co-application of D-AP5, and extracellular Ca^2+^/Na^+^-free medium with Na^+^ substituted by N-methyl-D-glucamine chloride (NMDG-Cl). Data were quantified from 3 experiments. **d**, Simultaneous imaging of mitochondrial Ca^2+^ using mito-GCaMP5G and mitochondrial membrane potential using TMRE during cLTD, with or without Ru360 (5 µM) to inhibit mitochondrial Ca^2+^ uptake. Data were quantified from 5 experiments for NMDA and 3 experiments for NMDA + Ru360. **e**, Measurement of mitochondrial membrane potential in isolated mitochondria following addition of 25 mM Na^+^. Data were quantified from 3 experiments. **f**, Simultaneous imaging of mitochondrial Ca^2+^ and membrane potential during cLTD with co-application of 5-(N-ethyl-N-isopropyl)amiloride (EIPA, 25 µM) to inhibit Na^+^/proton exchange activity. Data were quantified from 3 experiments. **g**, Quantification of TMRE fluorescence 5 min after NMDA addition from experiments shown in d and f. Data were quantified from 5 experiments for NMDA and 3 experiments for NMDA + EIPA. Statistical analysis was performed using an unpaired two-tailed t-test.

As we had observed for mitochondrial fission (Fig. 1e), cLTD-induced mitochondrial depolarization persisted even when extracellular Ca^2+^ was removed (Fig. 2c, green trace). Instead, mitochondrial depolarization was abolished only when both extracellular Ca^2+^ and Na^+^ were substituted with N-methyl-D-glucamine chloride (NMDG-Cl) to maintain osmolarity, suggesting that NMDA receptor-mediated Na^+^ influx is the major driver of cLTD-induced mitochondrial depolarization (Fig. 2c, red trace). To further validate that mitochondrial depolarization is uncoupled from mitochondrial Ca^2+^ uptake, we simultaneously monitored mitochondrial Ca^2+^ levels and mitochondrial membrane potential. Dual imaging revealed that mitochondrial Ca^2+^ levels increased transiently upon cLTD induction with a concomitant loss of membrane potential (Fig. 2d). However, inhibition of mitochondrial Ca^2+^ uptake with 5 µM Ru360 blocked mitochondrial Ca^2+^ accumulation without preventing mitochondrial depolarization (Fig. 2d), confirming that Ca^2+^was not the driving force for mitochondrial depolarization.

The mitochondrial membrane potential is generated through pumping of protons across the inner mitochondrial membrane. It was recently reported that Na^+^ ions contribute significantly to the mitochondrial membrane potential in mammalian mitochondria due to the action of the Na^+^/proton exchanger activity of mitochondrial complex I (*25*). To directly test whether the increase in cytosolic Na^+^ concentration commonly observed during cLTD (*26*) can drive mitochondrial depolarization, we treated isolated neuronal mitochondria with 25 mM Na^+^ and monitored their TMRE staining. Intriguingly, the addition of 25 mM Na^+^ triggered a loss in membrane potential (Fig. 2e). Consistently, inhibition of the complex I Na^+^/proton exchanger activity with 5-(N-ethyl-N-isopropyl)amiloride (EIPA) abolished cLTD-induced mitochondrial depolarization in intact neurons while preserving mitochondrial Ca^2+^ influx (Fig. 2f). Quantification of normalized TMRE fluorescence 5 min after NMDA addition confirmed that EIPA significantly attenuated mitochondrial depolarization compared with NMDA alone (Fig. 2g). Together, these results identify Na^+^-dependent mitochondrial depolarization as an upstream event during cLTD that occurs independently of cellular or mitochondrial Ca^2+^ uptake.

**Supplementary Fig. 2.**
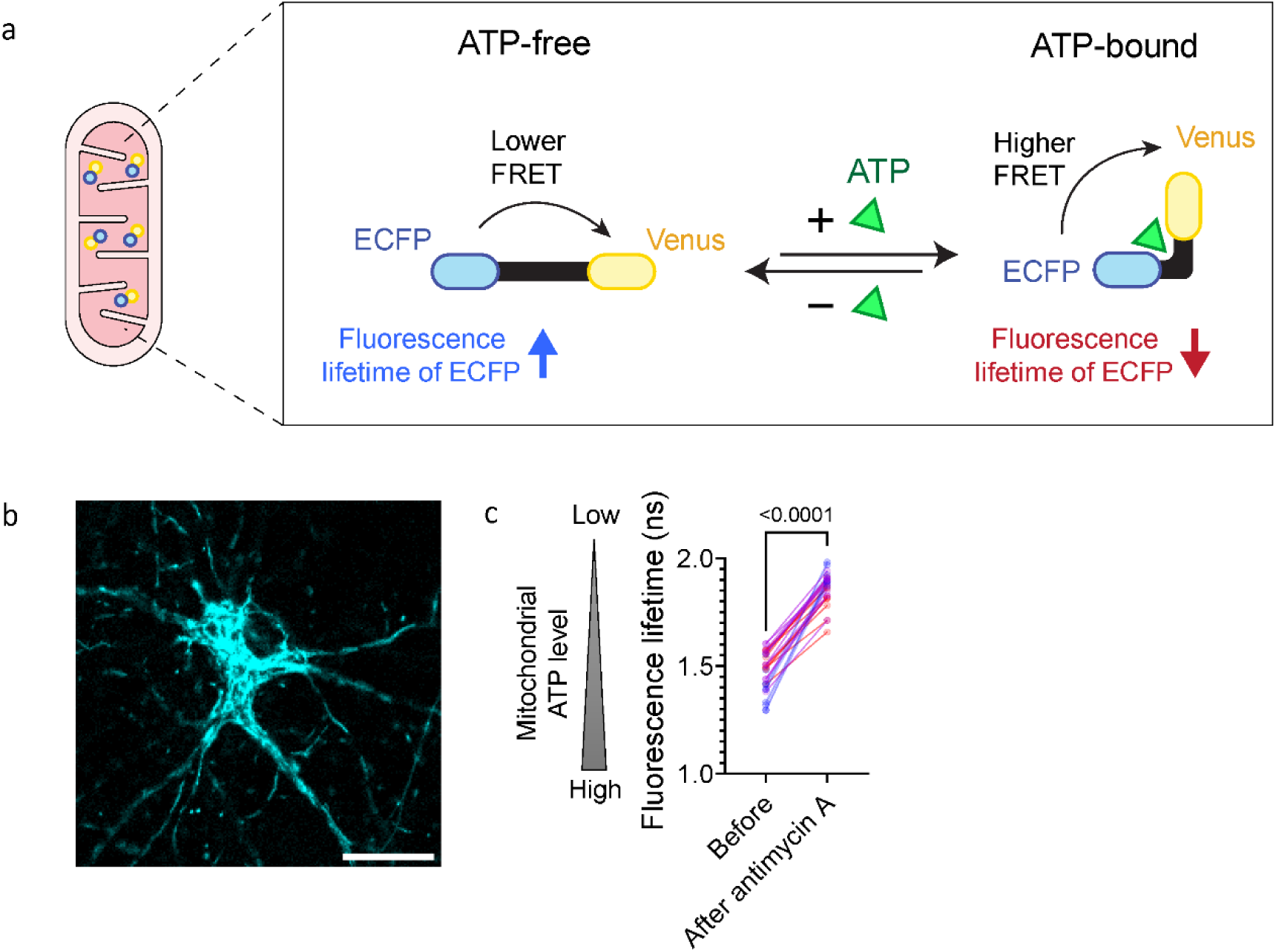
Mitochondrial ATP levels measured using the mito-ATeam ATP sensor. **a**, Schematic diagram of the mitochondrial-localized mito-ATeam sensor used to monitor ATP levels in mitochondria, and its molecular mechanism. **b**, Representative image of a neuron expressing mito-ATeam. Scale bar: 20µm. **c**, Calibration of mito-ATeam fluorescence lifetime after inhibition of mitochondrial ATP production with antimycin A (20µM). Data were quantified from 30 neurons from 3 experiments. Each point represents one neuron. Statistical analysis was performed using paired Student’s t-test.

### PINK1 synthesis is required for sustained mitochondrial fission during cLTD

Mitochondrial depolarization is commonly sensed by stabilization of the kinase PINK1 (*27*) and activation of PINK1 signaling has been linked to increased mitochondrial fission and mitophagy (*28–30*). However, the presence of this short-lived mitochondrial protein depends on protein translation (*31*, *32*), especially in distal neurites (*17*). Acute inhibition of translation with cycloheximide (CHX) prior to cLTD induction significantly reduced mitochondrial fission (Fig. 3a, white arrowheads and supplementary movie 2). This reduction in fission rate was also reflected in a lack of mitochondrial downscaling in dendrites by CHX treatment (Fig. 3b), indicating that ongoing protein synthesis is required for cLTD-associated mitochondrial remodeling. It is interesting to note that an initial, albeit transient, decrease in mitochondrial occupancy still occurs upon cLTD induction in the presence of CHX, which may correlate with mitochondrial swelling due to Na^+^ influx into the matrix (Fig. 3a, yellow arrowheads). To identify proteins preferentially translated under these conditions, we performed Translating Ribosome Affinity Purification (TRAP) followed by RNA sequencing (TRAP-seq) (*33*). GAGE (Generally Applicable Gene-set Enrichment) analysis of ribosome-associated transcripts revealed coordinated translation of mitochondrial pathways, including mitophagy, PINK1–PRKN-mediated mitophagy, and the intrinsic apoptotic pathway (Supplementary Fig. 3). Notably, cLTD increased ribosome association of transcripts encoding rapidly turned-over regulators of mitochondrial quality control and permeability, such as LC3A (an ATG8-family protein encoded by Map1lc3a gene), PINK1 and BOK (Fig. 3c). Among these candidates, PINK1 was of particular interest as we have previously shown that its mRNA is present in dendrites and axons and requires local translation for its activity (*17*). Intriguingly, mitochondrial depolarization promotes its stabilization and activation (*18*). This positions PINK1 as a potential molecular link between the translation dependence and mitochondrial depolarization underlying cLTD-induced mitochondrial remodeling.

**Fig. 3.**
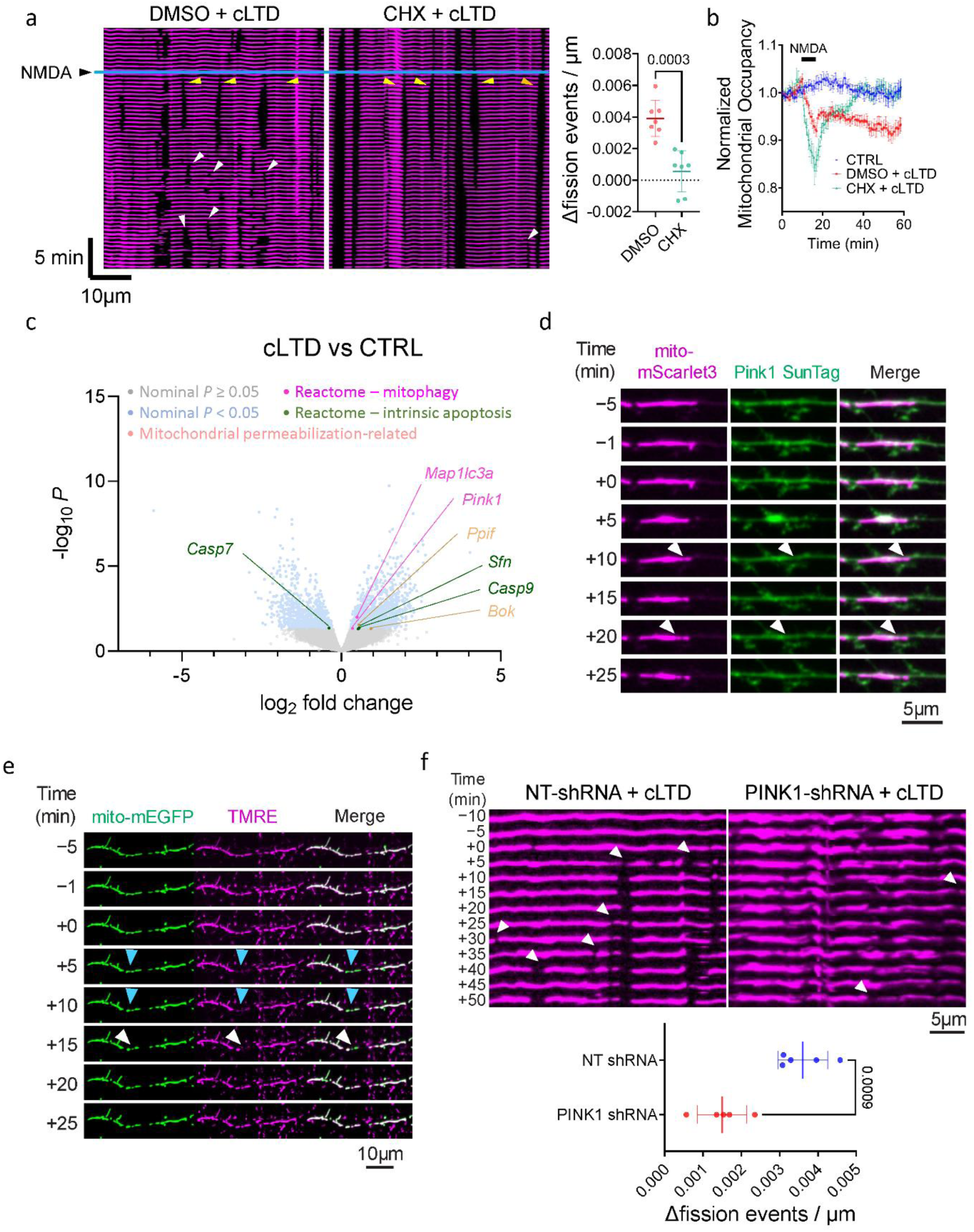
Local translation of PINK1 mediates mitochondrial remodeling downstream of Na^+^-dependent depolarization. **a**, Representative kymographs and quantification of mitochondrial fission events in dendrites following cLTD in the presence or absence of the translation inhibitor cycloheximide (CHX). Mitochondrial fission events were quantified from 7 experiments. Each point represents one neuron. Statistical analysis was performed using unpaired Student’s t-test. White arrowheads indicate mitochondrial fission events. Yellow arrowheads indicate mitochondrial swelling. See also supplementary movie 2. **b**, Quantification of normalized mitochondrial occupancy over time under control, cLTD, and cLTD + CHX conditions in Ca^2+^-free medium. Data were quantified from 7 experiments and presented as mean ± SEM. **c**, Volcano plot of differential abundance of ribosome-associated transcripts between cLTD and control conditions. Blue dots indicate genes with nominal *P* < 0.05, while grey dots indicate non-significant genes. Among genes with nominal *P* < 0.05, those annotated to Reactome mitophagy pathways (magenta) or the Reactome intrinsic apoptosis pathway (green), or literature-curated genes related to mitochondrial permeabilization (orange) are highlighted. **d**, Visualization of PINK1 using the SunTag system upon cLTD stimulation. A PINK1 construct fused to 10 tandem GCN4 epitopes was detected by co-expression with scFv-GCN4-GFP. White arrowheads indicate mitochondrial fission events. **e**, Representative time-lapse images of a dendritic segment from a DIV7 mouse primary hippocampal neuron expressing mito-mEGFP and labeled with TMRE. cLTD was induced at 0 min by NMDA application. mito-mEGFP marks mitochondrial morphology, whereas TMRE reports mitochondrial membrane potential. Arrowheads indicate a mitochondrion that undergoes loss of TMRE signal (depolarization, blue arrowhead) before subsequent fission (white arrowhead). Scale bar, 10 µm. See also supplementary movie 3. **f**, Representative kymographs and quantification of mitochondrial fission events during cLTD in neurons with PINK1 knockdown. Mitochondrial fission events were quantified from 5 experiments. Each point represents one neuron. Statistical analysis was performed using unpaired Student’s t-test. White arrowheads indicate mitochondrial fission events.

In order to directly visualize the stabilization of PINK1 on depolarized mitochondria, we employed a cytosolic PINK1 sensor that we previously developed (Pink1-ST-cyto (*34*)). This approach enables real-time detection of newly synthesized, cytoplasmic PINK1 by tagging PINK1 with tandem SunTag epitopes, which are recognized by an intracellularly expressed GFP-tagged single-chain antibody fragment (Supplementary Fig. 4). Using this system, we observed that PINK1 accumulated on depolarized mitochondria during cLTD (Fig. 3d), linking changes in mitochondrial membrane potential to spatially restricted PINK1 accumulation.

We next determined if mitochondrial depolarization precedes mitochondrial fission. NMDA application induced mitochondrial depolarization (blue arrowheads) before subsequent mitochondrial fission (white arrowheads) (Supplementary Fig. 5). Occasionally, depending on the synaptic maturity of the cultured neurons, this response was even limited to a single mitochondrion, which then proceeded to fission (Fig. 3e and supplementary movie 3). One part of this mitochondrion repolarized more quickly than the others (already white in the overlay of TMRE and mitoGFP at +15 min), suggesting that the matrix compartments are completely separated at this point.

The presence of PINK1 is required for mitochondrial fission during cLTD, as PINK1 depletion by shRNA significantly reduced mitochondrial fission during cLTD (Fig. 3f). Together, these results support a pathway in which mitochondrial depolarization during cLTD drives PINK1 activation to achieve sustained mitochondrial remodeling.

**Supplementary Fig. 3.**
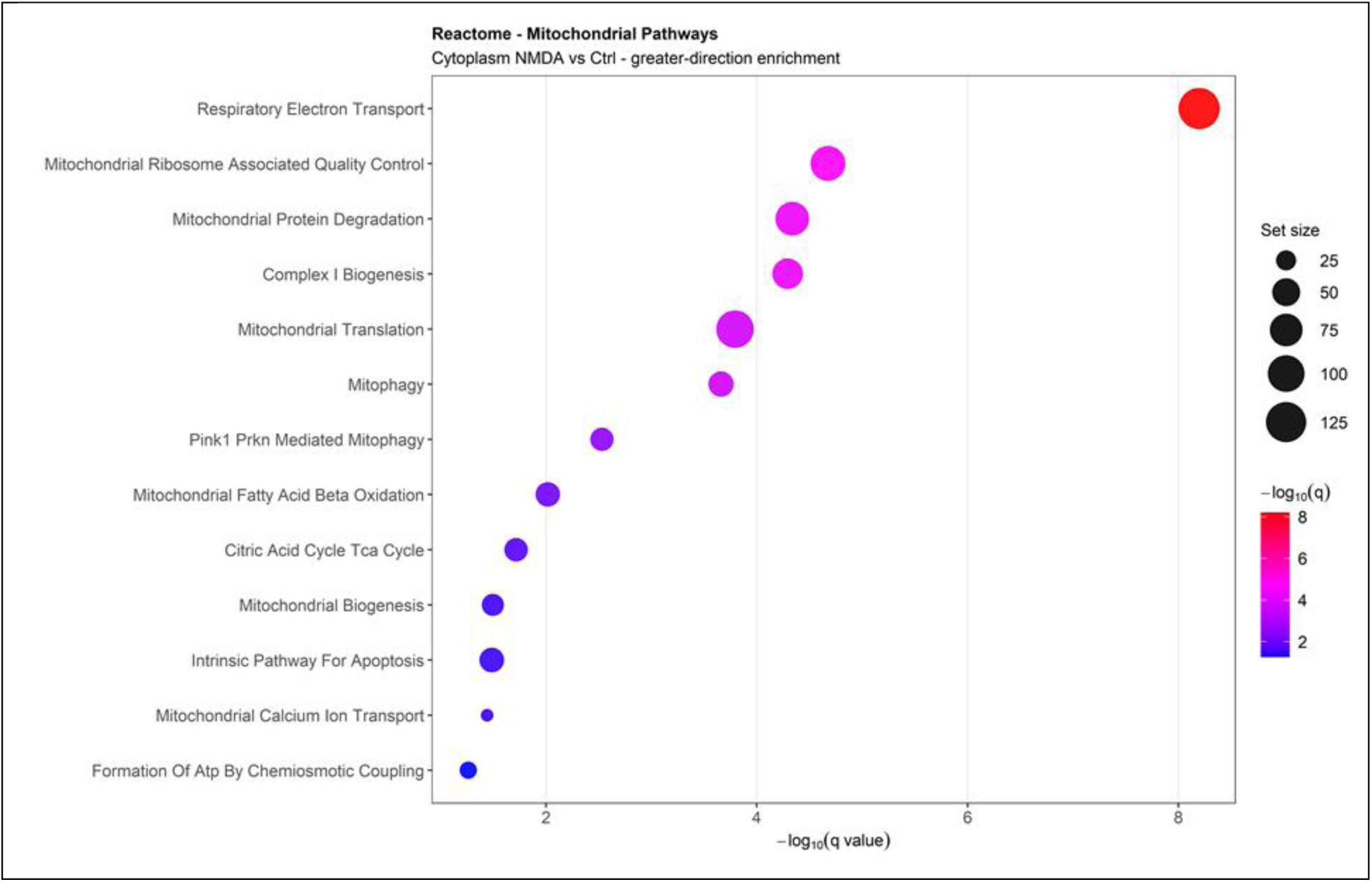
cLTD induces coordinated enrichment of mitochondrial pathways among ribosome-associated transcripts. GAGE (Generally Applicable Gene-set Enrichment) analysis of Reactome pathways using ribosome-associated transcripts cLTD-treated versus control neurons. Selected mitochondrial pathways showing greater-direction enrichment following cLTD are depicted. Dot position and color indicate −log10(q value), and dot size indicates pathway gene-set size.

**Supplementary Fig. 4.**
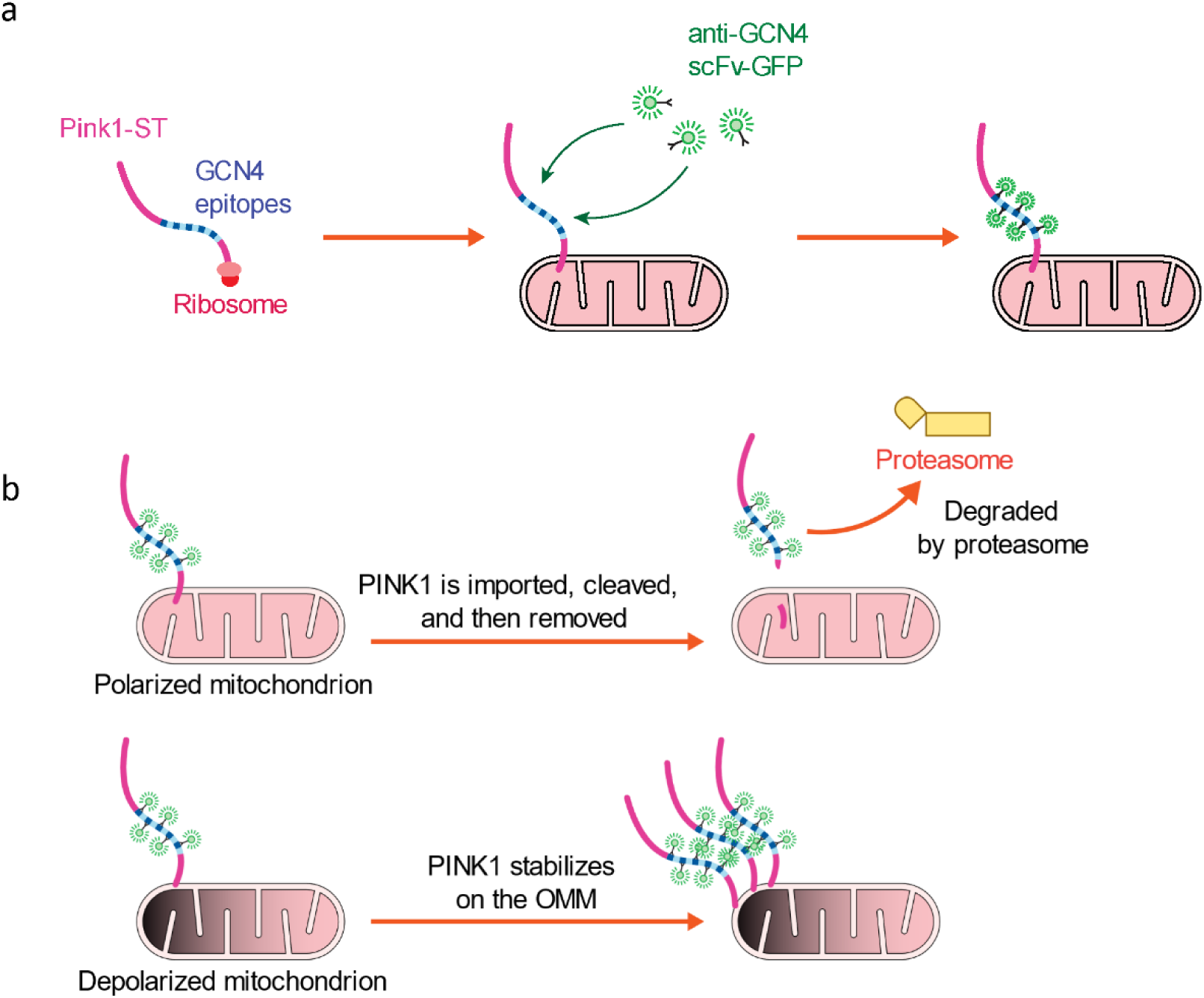
Schematic diagram of the SunTag-based reporter for newly synthesized PINK1. **a**, Schematic diagram showing PINK1 K218M kinase-dead–10×GCN4 reporter expressed together with scFv-GCN4-GFP. As newly translated PINK1 emerges, the ten GCN4 repeats recruit multiple scFv-GCN4-GFP molecules, thereby amplifying the GFP signal locally and enabling detection of newly synthesized PINK1. **b**, Under polarized mitochondrial conditions, newly synthesized PINK1 is imported into mitochondria, cleaved, and removed, followed by proteasomal degradation. Upon mitochondrial depolarization, PINK1 import is impaired, leading to stabilization of PINK1 on the outer mitochondrial membrane and accumulation of the SunTag signal.

**Supplementary Fig. 5.**
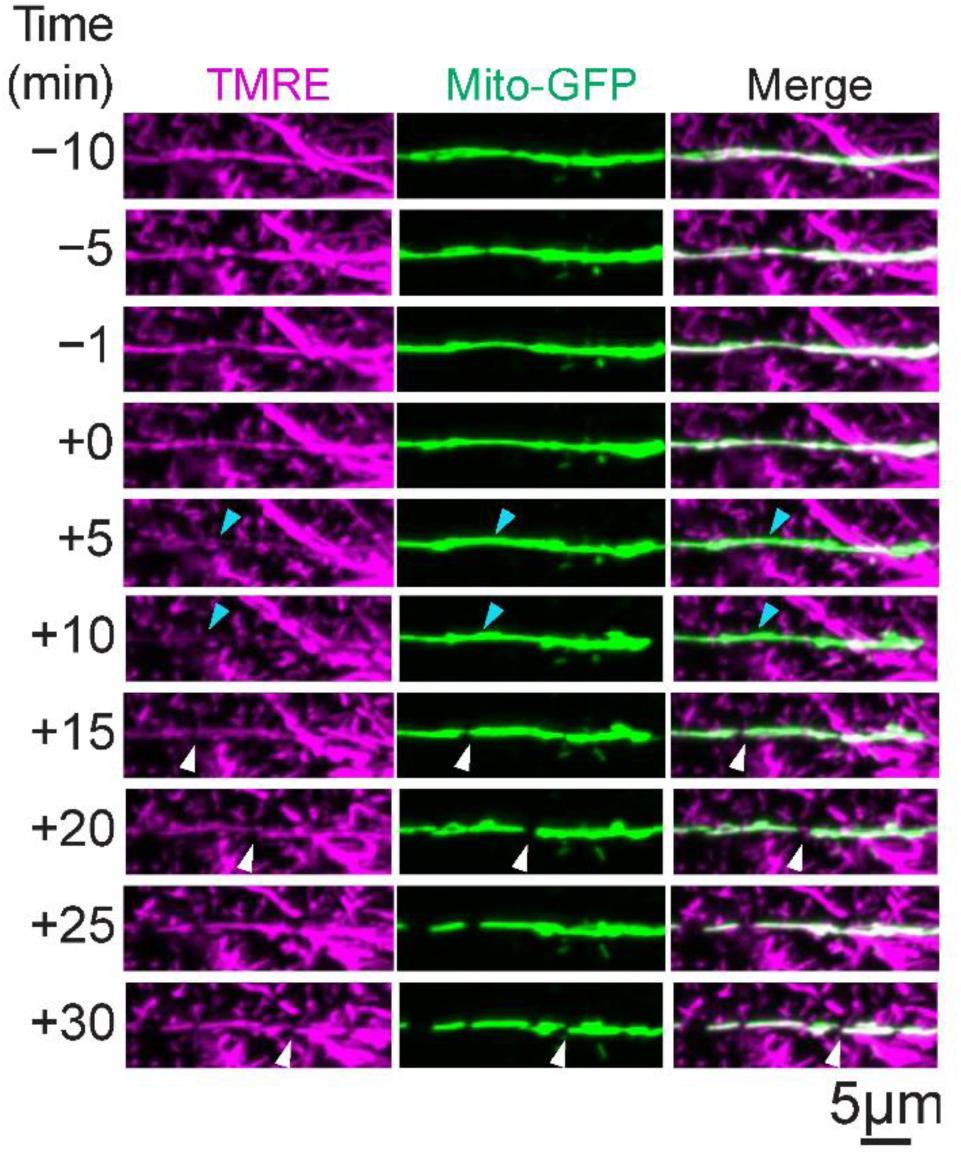
Mitochondrial depolarization precedes fission during cLTD. Representative time-lapse images of mitochondrial membrane potential using TMRE in hippocampal neurons expressing mito-GFP. cLTD was induced by NMDA application. Blue arrowheads indicate mitochondrial depolarization. White arrowheads indicate mitochondrial fission events.

### Na^+^-PINK1 signaling drives caspase-3 activation and spine remodeling during cLTD

Induction of cLTD has been previously linked to sub-lethal activation of caspase-3 (*9*) and mitochondrial fission has been shown to facilitate cytochrome c release and mobilization in apoptotic contexts (*35*, *36*). We therefore tested whether cLTD-associated mitochondrial remodeling engages caspase-3 signaling by targeting a caspase-3 cleavable FRET sensor (mCerulean-DEVD-mVenus; Fig. 4a, b; based on Rehm et al. (*37*)) to the mitochondrial outer membrane by addition of a CISD1 transmembrane anchor. Fluorescence lifetime-based imaging of the changes in donor lifetime (FLIM-FRET (*24*), validated by application of procaspase-activating compound PAC-1; Supplementary Fig. 6a, b) indicated increased cleavage of the FRET sensor, consistent with caspase-3 activation upon cLTD (Fig. 4c). The increase in caspase-3 cleavage could be blocked by the caspase-3 inhibitor Z-DEVD-FMK (Fig. 4d) and was dependent on ion influx through the NMDA receptor (Supplementary Fig. 6c). Inhibition of mitochondrial depolarization by EIPA was sufficient to prevent caspase-3 activation (Fig. 4e). Likewise, co-application of EIPA during cLTD also blocked spine shrinkage (Fig. 4f). This effect was dependent on the presence of PINK1, as cLTD-induced spine shrinkage was impaired in neurons with PINK1 knockdown (Fig. 4g). Together, these results identify a molecular pathway through which mitochondrial remodeling contributes to LTD. In this pathway, influx of Na^+^ through NMDA receptors triggers mitochondrial depolarization via dissipation of the Na^+^ gradient across the inner mitochondrial membrane, which is responsible for approximately half of the mitochondrial membrane potential (*25*). Depolarized mitochondria then accumulate newly synthesized PINK1, which promotes mitochondrial fission and engages downstream caspase-3 signaling. This mitochondrial remodeling pathway ultimately contributes to dendritic spine shrinkage during cLTD.

**Fig. 4.**
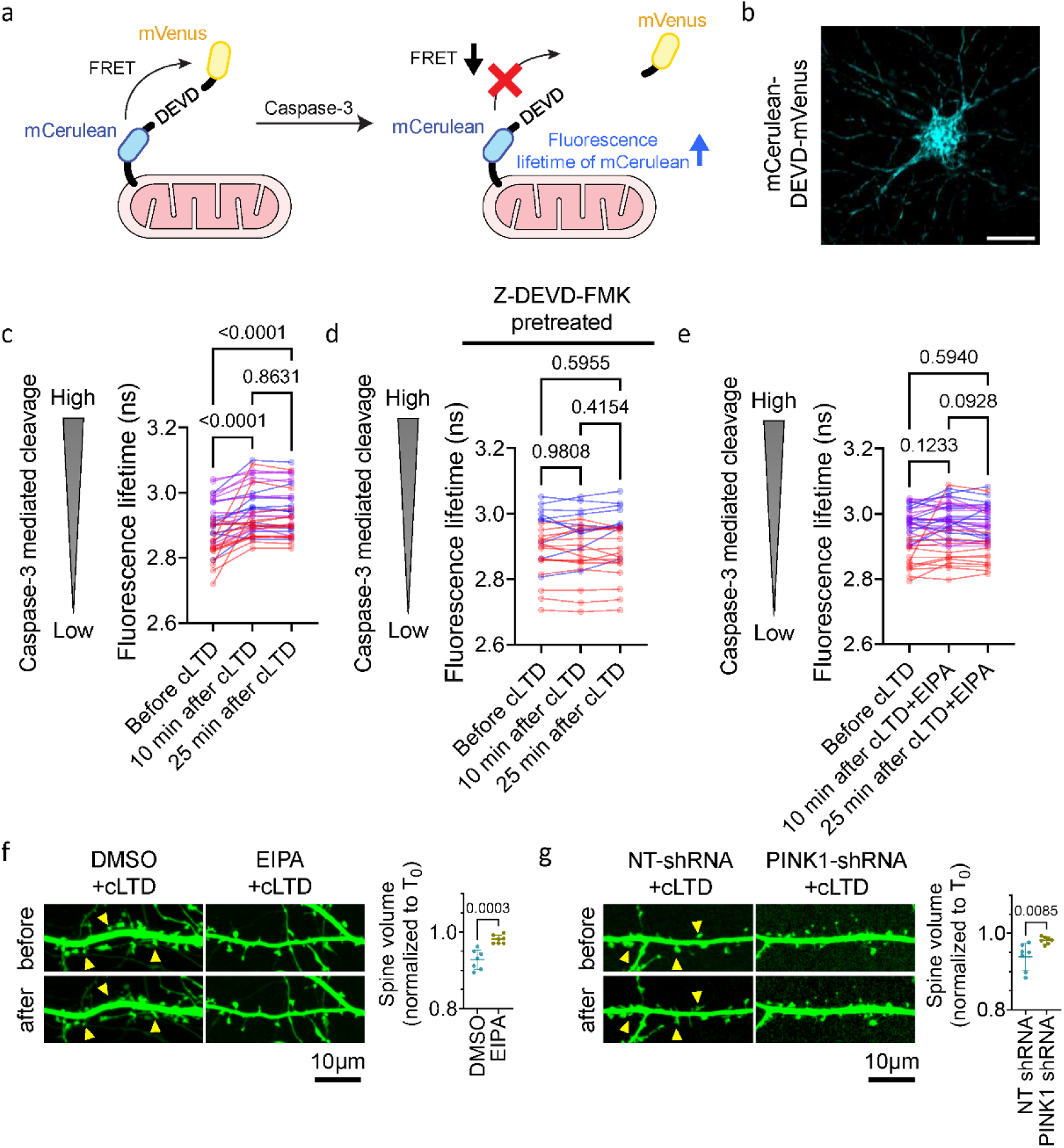
Na^+^–PINK1 signaling is required for caspase-3 activation and spine shrinkage during cLTD. **a**, Schematic of the FRET-based caspase-3 activity sensor CISD1-mCerulean-DEVD-mVenus. Cleavage of the DEVD linker by active caspase-3 reduces FRET efficiency, thereby increasing the donor fluorescence lifetime. **b**, Representative image of a neuron expressing CISD1-mCerulean-DEVD-mVenus. Scale bar: 20µm **c**, Quantification of caspase-3 activity after cLTD induction. Caspase-3 activity was quantified from 37 neurons from 3 experiments. Each point represents one neuron. Statistical analysis was performed using repeated-measures one-way ANOVA followed by Tukey post hoc tests. **d**, Quantification of caspase-3 activity after cLTD induction with Z-DEVD-FMK pretreatment. Caspase-3 activity following Z-DEVD-FMK pretreatment was quantified from 27 neurons from 2 experiments. Each point represents one neuron. Statistical analysis was performed using repeated-measures one-way ANOVA followed by Tukey post hoc tests. **e**, Quantification of caspase-3 activity during cLTD with co-application of 5-(N-ethyl-N-isopropyl)amiloride (EIPA) to inhibit Na^+^/proton exchange activity. Caspase-3 activity following EIPA treatment was quantified from 40 neurons from 3 experiments. Each point represents one neuron. Statistical analysis was performed using repeated-measures one-way ANOVA followed by Tukey post hoc tests. **f**, Representative image and quantification of dendritic spine size changes following cLTD in neurons treated with EIPA. Dendritic spine volume was quantified from 7 experiments. Each point represents one neuron. Statistical analysis was performed using unpaired Student’s t-test. Yellow arrowheads indicate dendritic spines undergoing morphological changes. **g**, Representative image and quantification of dendritic spine size changes following cLTD in control and PINK1 knockdown neurons. Dendritic spine volume was quantified from 7 experiments. Each point represents one neuron. Statistical analysis was performed using unpaired Student’s t-test. Yellow arrowheads indicate dendritic spines undergoing morphological changes.

**Supplementary Fig. 6.**
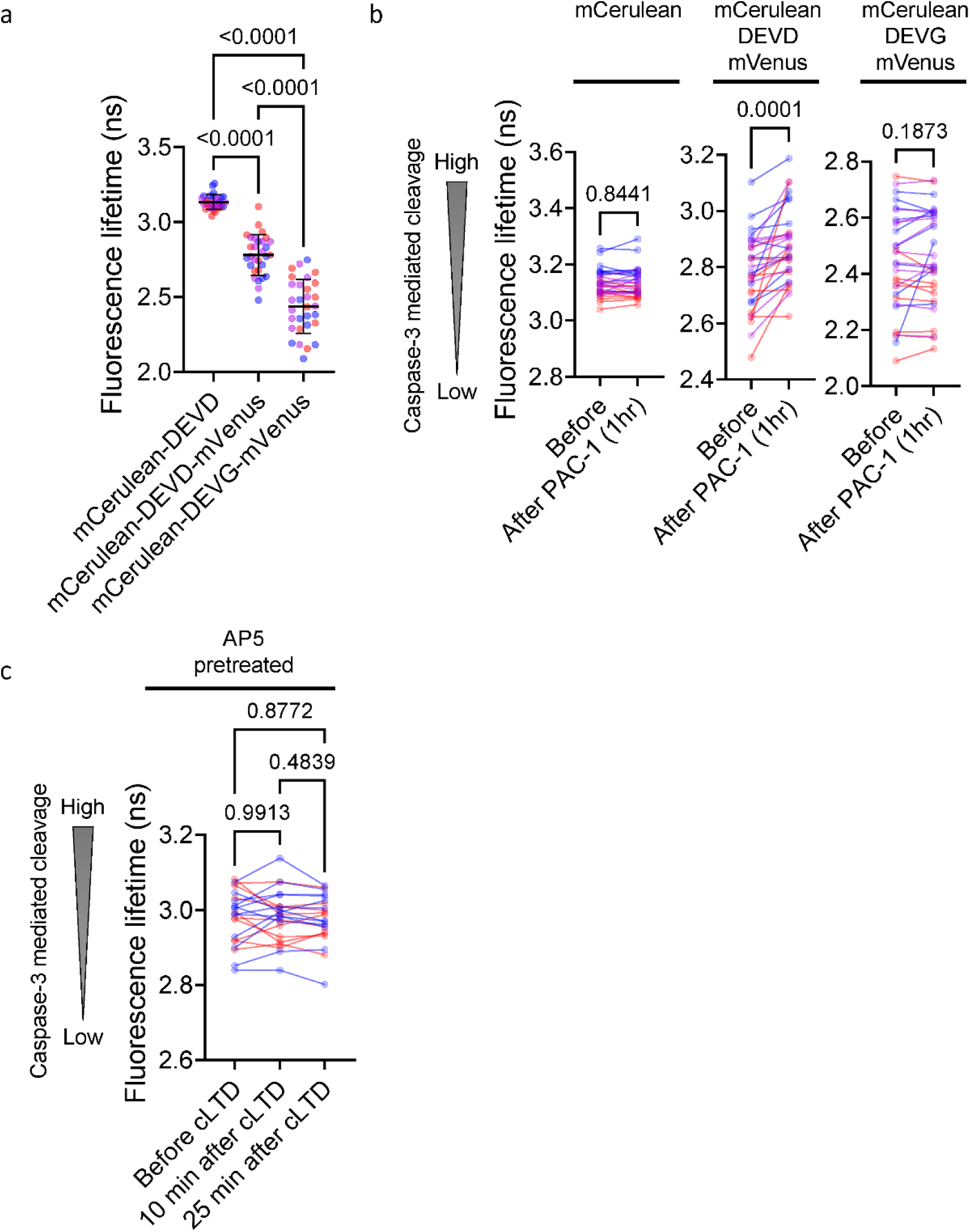
Validation of the FLIM-based mitochondrial targeted caspase-3 activity sensor. **a**, Fluorescence lifetime measurements of control constructs containing CISD1-mCerulean, CISD1-mCerulean fused to the caspase-3-cleavable DEVD linker and mVenus, or CISD1-mCerulean fused to the non-cleavable control linker DEVG and mVenus. Donor fluorescence lifetime was quantified from 30 neurons from 3 experiments. Each point represents one neuron. Statistical analysis was performed using one-way ANOVA followed by Tukey post hoc tests. **b**, Validation of caspase-3-dependent sensor cleavage using the procaspase-activating compound PAC-1. Donor fluorescence lifetime before and after PAC-1 treatment was quantified for the cleavable mCerulean-DEVD-mVenus sensor, mCerulean-DEVD donor-only control, and non-cleavable mCerulean-DEVG-mVenus control. Data were quantified from 30 neurons from 3 experiments. Each point represents one neuron. Statistical analysis was performed using paired Student’s t-test. **c**, Donor fluorescence lifetime of the DEVD sensor in D-AP5-pretreated neurons before and after cLTD induction. Donor fluorescence lifetime was quantified from 24 neurons from 2 experiments. Each point represents one neuron. Statistical analysis was performed using repeated-measures one-way ANOVA followed by Tukey post hoc tests.

## Discussion

Here we show that activation of PINK1 is a defining feature of cLTD and necessary for mitochondrial fission and spine changes. In contrast to LTP, where mitochondrial changes are driven by Ca^2+^-dependent activation of CamKII, the main driver of the mitochondrial response is the influx of Na^+^ ions through NMDA receptors. This fits well with the reduced influx of Ca^2+^ during cLTD which is not sufficient to drive CamKII activation to comparable levels as during LTP (*38*, *39*). Instead, Na^+^-influx dependent mitochondrial depolarization and PINK1 activation are required to initiate fission (Fig. 3). Increased fission, along with Na^+^-induced swelling, facilitates the sub-lethal activation of caspase-3, as mitochondrial outer membrane permeabilization (MOMP) is more easily observed in smaller mitochondria with higher membrane curvature (*40*, *41*). The transient nature of mitochondrial depolarization and the delayed fission due to the necessity to translate new proteins, including PINK1 and other apoptotic regulators (Fig. 3c), may be another key to restrict MOMP to a subset of mitochondria to prevent widespread MOMP and excitotoxic apoptosis induction. This enables the non-apoptotic activation of caspase-3 (Fig. 4) necessary for structural changes to spines (*42–44*). Furthermore, PINK1/Parkin-dependent mitophagy or MDV generation can explain the small but persistent reduction in mitochondrial occupancy we observe upon cLTD (Fig. 1c and 3b). This fits with the observed role for autophagic processes within dendrites that support the progression of LTD (*45*).

The discovery of transient mitochondrial depolarization and PINK1 activation as recipients of NMDA-driven Na^+^ influx sheds new light on the role of Na^+^ ions as second messengers in neurons. Na^+^ influx has been proposed to provide positive feedback on NMDA receptor activity as well as to alter mitochondrial Ca^2+^ content via the Na^+^/Ca^2+^ exchanger NCLX (*46*, *47*). Indeed, we observe that the clearance of Ca^2+^ from mitochondria is altered upon inhibition of the NHE function of complex I (Fig. 2d vs 2f). The importance of a separate Na^+^ gradient for the maintenance of the mitochondrial membrane potential has only recently been appreciated (*25*). Mechanistically, the NHE function of complex I converts part of the proton gradient into a Na^+^ gradient, which slows the dissipation of the proton gradient through the activity of complex V. Influx of Na^+^ through activation of the NMDA receptor transiently increases the cytosolic Na^+^ concentration, which is transmitted through the voltage-gated anion channel (VDAC) into the intermembrane space (IMS) of mitochondria. This triggers the reverse action of the NHE function of complex I leading to the electro-neutral exchange of one proton for each Na^+^ ion. In one possible model, this additional acidification of the IMS is used up by the F_1_F_O_-ATP-Syntase (complex V). This would predict an initial hyperpolarization as is observed in pancreatic beta cells (*47*), however, we do not record such an event. This may either be due to the fast nature of ion exchange (below our imaging rate of 30 seconds per frame), or it may suggest the existence of additional uncoupling mechanisms triggered by Na^+^ influx in neuronal mitochondria. Given the tight buffering and much lower intracellular concentration of free Ca^2+^ compared with Na^+^, Na^+^ influx may impose a greater osmotic burden on mitochondria during cLTD. We propose that mitochondrial uncoupling may be a consequence of the Na^+^-induced swelling (Fig. 3a, yellow arrowheads) and accompanied by a transient permeabilization of the inner membrane. Such a phenomenon can be observed in a neuronal cell line undergoing apoptosis (*48*) and thus may be tied to a rearrangement of the IMM and cytochrome c release for caspase-3 activation. In the case of LTD-induced Na^+^ influx, however, both mitochondrial swelling and depolarization are reversible, thus creating a sub-lethal stimulus to allow local caspase-3 signaling without cell death. This occurs even in the absence of Ca^2+^ influx into mitochondria (Fig. 2d), ruling out Ca^2+^-overload triggered permeability transition. This contrasts with excitotoxic cell death, which can be elicited by high-dose NMDA treatment (200 µM) and depends on mitochondrial Ca^2+^ overload (*49*).

Mitochondrial depolarization and ATP loss can be inhibited by the import of cytoplasmic ATP into mitochondria via the ADP/ATP carrier. This supports the reverse proton flow via the F_1_F_O_-ATP-Syntase (*50*), which turns into an ATPase to pump protons out of the matrix to quickly restore membrane potential. However, neuronal mitochondria express high levels of the ATPase inhibitory factor 1 (ATPIF1) that prevents the reverse action of the ATPase (*51*, *52*). This may cause the observed drop in mitochondrial ATP levels (Fig. 2a), preventing a detrimental depletion of cytoplasmic ATP levels (*51*, *52*). Importantly, ATPIF1 has been identified as an essential factor that allows sufficient time for PINK1 activation (*53*), setting the stage for PINK1 signaling.

Synaptic plasticity is thought to be the cellular equivalent of learning and memory. Mutations for PINK1 are found in hereditary forms of Parkinson’s disease (PD) (*54*), which can include cognitive and psychiatric symptoms in addition to parkinsonism (*55*). While PINK1 knockout mice fail to reproduce many motor aspects of the disease, it was recently reported that they display learning deficits (*56*), supporting a role for PINK1 in memory formation. Similar cognitive impairments have also been observed in mice lacking the PINK1 interactor Parkin (*57*), suggesting that Parkin may similarily contribute to synaptic plasticity similarly. Together, PINK1 and Parkin have been best studied as initiators of damage-induced mitophagy (*58*). However, despite abundant evidence for the importance of PINK1 for the detection of depolarized mitochondria in cell culture, the amount of PINK1-dependent mitophagy in neuronal cell bodies *in vivo* is negligible under baseline conditions (*59*). Our results now reveal a physiological stimulus for PINK1 during cLTD that can explain the importance of this protein for neurons beyond its role in mitophagy.

## Materials and methods

### Animals and primary hippocampal neuron culture

Primary hippocampal neurons were prepared from embryonic day 16.5 (E16.5) C57BL/6NRj wild-type mouse embryos as previously described (*17*). All animal procedures were approved by the Government of Upper Bavaria and performed in accordance with animal ethics guidelines. Briefly, timed-pregnant mice were euthanized by CO_2_ inhalation, embryos were rapidly isolated, and hippocampi were dissected and collected in ice-cold dissociation medium consisting of Ca^2+^-free Hank’s Balanced Salt Solution (ThermoFisher Scientific) supplemented with 10 mM MgCl₂, 1 mM kynurenic acid (Sigma-Aldrich), and 10 mM HEPES. Hippocampal tissue was enzymatically dissociated with papain/L-cysteine (Sigma-Aldrich) for 5 min at 37 °C, followed by trypsin inhibitor treatment and gentle mechanical trituration. Dissociated neurons were plated on acid-washed glass coverslips or glass-bottom plates (CellVis) coated with poly-L-lysine (Sigma-Aldrich) and laminin (ThermoFisher Scientific), and maintained in Neurobasal medium supplemented with B27 (2%, Gibco), L-glutamine (2 mM, Gibco), and penicillin/streptomycin (100 units, Gibco). Unless otherwise indicated, experiments were performed in hippocampal neurons at DIV14.

### DNA constructs, biosensors and neuronal transfection

Neurons were transfected with plasmids encoding fluorescent reporters, biosensors or shRNA using Lipofectamine 2000. For neurons grown in 24-well glass-bottom plates, 0.3 µg plasmid DNA was mixed with 1 µl Lipofectamine 2000 per well to prepare the transfection mixture. Before transfection, conditioned culture medium was replaced with B27-free Neurobasal medium. The transfection mixture was added to the cells for 20 min, after which the medium was replaced with B27-containing conditioned medium.

For analysis of dendritic morphology and mitochondrial dynamics, neurons were transfected with cytosolic GFP and mitochondria-targeted mito-mScarlet. Mitochondrial morphology and membrane potential were visualized using mito-GFP or mito-mEGFP together with TMRE. Mitochondrial ATP levels were monitored with a mitochondria-localized ATeam FRET sensor. Mitochondrial Ca^2+^ was monitored using mito-GCaMP5G. Newly synthesized PINK1 was visualized using a SunTag-based reporter consisting of kinase-dead PINK1 K218M fused to 10xGCN4 epitopes (Pink1-ST-cyto, (*34*)) and co-expressed with scFv-GCN4-GFP. Caspase-3 activity at the mitochondrial outer membrane was monitored using the CISD1-mCerulean-DEVD-mVenus FRET sensor; donor-only and non-cleavable DEVG constructs were used as controls where indicated and were transfected 2 days prior to imaging.

For TRAP-seq analysis, the plasmids required for virus production pMDLg/pRRE, pMD2.G and pRSV-Rev were acquired from Addgene (#12251, #12259 and #12253, respectively). The Pink1-kinase dead-10xGCN4cyto-12xMS2-PP7 (Pink1-SunTag) was described before (*34*), by inserting 10x GCN4 consecutive repeats (obtained by enzymatic restriction) in frame between the amino acid positions 141-145 of rat PINK1-K218M mutant. The plasmid coding for scFv-GCN4-sfGFP was acquired from Addgene (# 60907). The cytosolic BirA-mCherry construct was reported before (*34*) and was generated by amplifying BirA-mCherry including the Tet-O inducible promoter from the Addgene plasmid #62366, including a stop codon by site directed mutagenesis by PCR. The PCR product was then inserted into the vector backbone pMK1047 of the Addgene plasmid #62365 using the restriction enzymes NheI and EcoRI. Constructs were grown on DH5α chemically competent *E*. *coli* bacteria (strain Invitrogen, prepared in-house).

For PINK1 knockdown, neurons were transfected with shRNA targeting mouse Pink1 (Sigma-Aldrich TRCN0000026743) or a non-targeting control shRNA (Sigma-Aldrich SHC016).

### Induction of chemical synaptic plasticity

Chemical long-term depression (cLTD) and chemical long-term potentiation (cLTP) were induced using protocols adapted from existing protocols (*6*, *60*). Unless specified, cLTD was induced by bath application of NMDA (20 µM) for 5 min in Hibernate-E imaging medium without phenol red (BrainBits), followed by washout into fresh imaging medium. For cLTP experiments, neurons were incubated with D-AP5 (20 µM) for 48 h before imaging. Before stimulation, cells were preincubated for 1 h in imaging medium containing D-AP5 (20 µM, Sigma-Aldrich), TTX (0.5 µM, Acros Organics), strychnine (1 µM,Carl Roth GmbH), and picrotoxin (50 µM, Sigma-Aldrich). Immediately before stimulation, the preincubation buffer was washed with Mg^2+^-free imaging buffer containing 20 µM bicuculline and 1 µM strychnine, and cLTP was induced by application of glycine (400 µM) for 15 min. After stimulation, cells were washed with Mg^2+^-containing imaging solution. Where indicated, NMDAR activation was blocked by pretreatment or co-application of D-AP5 (40 µM). For experiments with Na^+^- and Ca^2+^-displaced medium, the following medium is used: NaCl (150 mM), KCl (5 mM), CaCl_2_ (2 mM), MgCl_2_ (2 mM), HEPES (10 mM, pH 7.4), D-Glucose (30 mM). An equal molar of Na^+^ or Ca^2+^ was substituted with N-methyl-D-glucamine chloride (NMDG-Cl) in the imaging medium. Pharmacological compounds were applied as follows: cycloheximide (CHX, 10 µg/ml, Enzo Life Sciences), Ru360 (5 µM, Sigma-Aldrich), 5-(N-ethyl-N-isopropyl)-amiloride (EIPA, 25 µM, Sigma-Aldrich), Z-DEVD-FMK (5 µM, Tocris Bioscience), antimycin A (20 µM, Sigma-Aldrich), and PAC-1 (20 µM, Adooq Bioscience).

### Live-cell fluorescence imaging

Live-cell imaging was performed at the Imaging Facility of the Max Planck Institute for Biological Intelligence, Martinsried, Germany. Images were acquired using a Nikon Eclipse Ti2 spinning-disk microscope equipped with a CSU-W1 SoRa spinning-disk confocal scanning unit (Yokogawa Electric), a 60x oil-immersion objective (Nikon), and a Photometrics Prime BSI sCMOS camera (Teledyne Vision Solutions). Image acquisition was controlled using NIS-Elements software version 5.21.03. Unless otherwise specified, cells were imaged at 37 °C in Hibernate-E imaging medium. For TMRE imaging, neurons were loaded with 10 nM TMRE in Hibernate-E at 37 °C for 30 min before imaging.

### Image analysis and quantification

Image analysis was performed using ImageJ. Time-lapse images were stabilized using the ImageJ image stabilizer plugin to correct unwanted x-y stage movements during acquisition (*61*). Images were corrected for photobleaching in ImageJ using an exponential decay correction calculated from the mean fluorescence intensity of the analyzed cell/neurite region after background subtraction. Dendritic regions of interest were selected from healthy, isolated dendritic segments and then straightened using the built-in ImageJ Straighten function for easier analysis and display. Mitochondrial occupancy was calculated as the total length occupied by mitochondria divided by the dendrite length and was normalized to the pre-stimulation baseline. Mitochondrial fission events were identified from straightened kymographs using criteria similar to previously described methods (*62*). Fission events were normalized to dendrite length and time. TMRE and mito-GCaMP fluorescence values were background-subtracted and normalized to baseline.

### FLIM-FRET imaging of mitochondrial ATP and caspase-3 activity

Fluorescence lifetime imaging microscopy (FLIM) was used to quantify changes in Förster resonance energy transfer (FRET) in donor-acceptor biosensors. FLIM imaging was performed at the Imaging Facility of the Max Planck Institute for Biological Intelligence, Martinsried, Germany, using a Leica SP8 confocal laser-scanning microscope equipped with an 63× water-immersion objective (Leica) and LAS X software version 3.5.5 with the FLIM module (Leica). Imaging was performed at 37 °C. A pulsed 440 nm laser (PicoQuant) was used to excite the donor fluorophore of each construct. Donor fluorescence lifetime was determined from photon arrival times using phasor analysis method (*63*).

For mito-ATeam measurements, we used a construct similar to one previously described (*23*). The sensor consists of the ATeam ATP FRET reporter fused at its N-terminus to two tandem repeats of the cytochrome c oxidase subunit VIII mitochondrial targeting sequence, which directs the reporter to the mitochondrial matrix. Lifetime changes of ECFP were used as a readout of mitochondrial ATP levels. Sensor validation was performed by inhibition of mitochondrial ATP production with antimycin A. For caspase-3 measurements, lifetime changes of mCerulean were monitored. Cleavage of the DEVD linker reduces FRET efficiency and increases donor fluorescence lifetime. Sensor specificity was assessed using donor-only, cleavable DEVD, and non-cleavable DEVG controls, PAC-1-induced procaspase activation, D-AP5 pretreatment, and Z-DEVD-FMK inhibition.

### Isolation of neuronal mitochondria and Na^+^ stimulation

Mitochondria were isolated from cultured neurons using a protocol adapted from (*64*). In brief, cells were homogenized in ice-cold buffer containing 220 mM mannitol, 70 mM sucrose, 1 mM EDTA, 0.5 mM PMSF, 2 mg/ml bovine serum albumin, and 20 mM HEPES-KOH, pH 7.6, and mitochondria were isolated by differential centrifugation. Isolated mitochondria were collected and resuspended in buffer containing 250 mM sucrose, 5 mM magnesium acetate, 80 mM potassium acetate, 10 mM sodium succinate, 1 mM dithiothreitol (DTT), 5 mM ATP, and 20 mM HEPES-KOH, pH 7.4. To test the direct effect of Na^+^ on mitochondrial membrane potential, 25 mM NaCl was acutely added, and TMRE fluorescence was monitored over time. Fluorescence values were normalized to the initial baseline intensity.

### Lentivirus production

Lentiviral particles were generated using HEK293T cells as previously described (*65*). The cells were seeded onto collagen type-I (Sigma-Aldrich)-coated 10 cm dishes (Falcon) at a density of 6×106 cells per dish and cultured in DMEM containing 10 % FBS and 1 % P/S. After 24 h, each 10 cm dish of HEK293T cells underwent transfection with 5 µg of the packaging plasmid mix (pMDLg/pRRE, pRSV-Rev, pMD2.G; ratio 4:1:1) and 5 µg of the relevant transfer plasmid (FudeltaGW-rtTA, AviTag RPL10a or cytosolic BirA-mCherry) using TransIT-Lenti reagent (Mirus Bio). After 48 h, the cell culture medium containing lentiviral particles was harvested, clarified by centrifugation at 300 x g for 10 min at room temperature, and then mixed with lentivirus precipitation solution (Alstem) at a 4:1 ratio. The mixture was incubated at 4 °C overnight before undergoing centrifugation at 1,500 x g for 30 min at 4 °C to precipitate lentiviral particles. After discarding the supernatant, the pellet containing lentiviral particles was reconstituted in 1 ml of ice-cold PBS per 10 cm dish, aliquoted and stored at -80 °C.

### Translating Ribosome Affinity Purification followed by RNA Sequencing (TRAP-seq)

RNA associated with translating ribosomes was affinity purified using an AviTag–BirA-based biotinylation system similar to previously described (*33*). In brief, mouse primary cortical neurons were seeded in 6-well plates (Greiner) coated with PLL and laminin at a density of 2×10^6^ cells per well. On DIV12, half of the culture media was collected and neurons were co-transduced for 5-8 h with AviTag-RPL10a, rtTA and BirA-mCherry lentiviral particles. After that, neurons were rinsed three times with wash medium (Neurobasal +P/S/G) and then maintained in NB medium (Neurobasal+ 1x B27 +P/S/G) supplemented 1:1 with the previously collected conditioned medium. On DIV14, neurons were treated with doxycycline (2 ng/µL) for 4 h to induce BirA-mCherry expression. Subsequently, neurons were stimulated with NMDA for 5 min in Hibernate-E medium to induce NMDA receptor-dependent long-term depression (LTD), while control neurons were incubated in Hibernate-E medium without NMDA. Following treatment, both NMDA-stimulated and control neurons were rinsed and incubated for an additional 10 min in Hibernate-E medium. Neurons were subsequently lysed in a buffer containing 25 mM Tris/HCl pH 7.4, 1 % NP-40, 150 mM NaCl, supplemented with 1X protease inhibitor cocktail (Roche), 200 µM PMSF, and 1% Murine RNase Inhibitor (NEBiolabs). After centrifugation at 9,300 x g for 1 min, the pellets were discarded and 10 % of the supernatant was kept as input, while the remaining cell lysate was incubated with 30 µl Streptavidin-Agarose beads (Sigma-Aldrich) per sample for 24 h at 4 °C. Afterwards, the beads were washed by centrifugation (3 minutes, 1,000 x g, 4 °C) four times in lysis buffer, as before. RNA isolation of both input and pulldowns was performed with the RNaeasy Kit (Qiagen) following manufactures instructions.

Library prep was performed from 10 ng RNA per sample using the Takara SMART-Seq® Total RNA Single Cell Library Prep with ZapR® Depletion (cat #634360). Sequencing was performed on Element Bioscience AVITI 2x 75bp, 9M reads per sample on average (ranging from 5M to 15M reads).

After quality control, reads were aligned to the mouse reference genome GRCm39 using the STAR aligner (*66*). Gene expression normalization and differential expression analysis were conducted using the limma package from Bioconductor (*67*, *68*). Genes were filtered to retain only those with a total count greater than 2 across all samples (i.e., a row sum > 2 in the gene-by-sample matrix). Normalization was performed using the trimmed mean of M-values (TMM) method, accounting for library size. Differentially expressed genes were identified based on adjusted *p*-values calculated using the Benjamini–Hochberg method, applying a significance threshold of *p* < 0.01.

Functional enrichment analysis was performed with the GAGE package, using the mouse MSigDB CP:REACTOME signaling pathways (*69*, *70*). Pathways with an adjusted p. value < 0.1 (Benjamini-Hochberg) were considered significantly enriched.

### Statistical analysis

Statistical analysis was performed using GraphPad Prism 10.6.1. Data are presented as mean ± SD unless otherwise stated. Each data point represents one neuron unless otherwise indicated. For paired before-after comparisons, paired Student’s t-tests were used. For comparisons between two independent groups, unpaired Student’s t-tests were used. For multiple-group comparisons, one-way ANOVA or repeated-measures one-way ANOVA was used as appropriate, followed by Tukey post hoc tests. Statistical significance was defined as p < 0.05.

## Acknowledgments

We thank J. Lindner for technical support and all the members of the Harbauer laboratory for their support and many fruitful discussions. We are grateful to R. Kasper/E. Laurell/C. Polisseni and M. Spitaler/M. Oster from the Imaging facilities of the MPIs for Biological Intelligence (RRID:SCR_026797) and Biochemistry (RRID:SCR_025739), respectively, for assistance with live cell imaging. We would also like to thank R. Kim from NGS Core Facility of the MPI of Biochemistry (RRID:SCR_025746)

## Funding

Work in ABH’s lab is supported by the Max Planck Society, the German research foundation (DFG, HA 7728/6-1 – ID 576693647, EXC 2145 SyNergy – ID 390857198, TRR353 – ID 471011418, SPP2453 ID 541742535), the European Union (ERC StG Project 101077138 — MitoPIP) and Germany’s Excellence Strategy within the framework of the Munich Cluster for Systems Neurology (EXC 2145 SyNergy – ID 390857198) and the Schram foundation (T0287/46550/2025). Views and opinions expressed are however those of the author(s) only and do not necessarily reflect those of the European Union or the European Research Council. Neither the European Union nor the granting authority can be held responsible for them.

Work in MR’s laboratory is supported by the Deutsche Forschungsgemeinschaft (DFG) under DFG grants INST 38/655-1 (ID 471011418) TRR 353, DFG GRK 3112/1 (ID 538201975), DFG grant MO 3226/4-1 and through Germany’s Excellence Strategy, DFG grant EXC 2075 (ID 390740016).

Work in MB’s lab is supported by the Deutsche Forschungsgemeinschaft (DFG) – TRR 353 (Project ID 471011418 - SP02), CRC1479 (Project ID 441891347 - S1), CRC 1160 (Project ID 256073931 - Z02), CRC1453 (Project ID 431984000 - S1), TRR167 (Project ID 259373024 - Z01), TRR417 (Project ID 540805631 - S03), FOR 5476 UcarE (Project ID 493802833-P7) and the German Federal Ministry of Research, Technology and Space (BMBFTR) – PM4Onco (FKZ 01ZZ2322A) and SATURN3 (01KD2206L)

## Author Contributions

ABH conceived the project and wrote the manuscript together with BCKT, with additional input from IS, SW, AM, MB and MR. BCKT designed and conducted most of the experiments. IS was responsible for the experiments using isolated mitochondria and TRAP-seq. SW and MR were responsible for the development of the caspase-3 sensor. AM and MB were responsible for the bioinformatic analysis. All authors read and approved the manuscript.

## Declaration of Interests

The authors declare no competing interests.

## List of Supplementary Materials

**Supplementary Table 1. GAGE (Generally Applicable Gene-set Enrichment) analysis of ribosome-associated transcripts profiled by TRAP-seq following cLTD.** Reactome pathways exhibiting coordinated increases or decreases in ribosome-associated transcript abundance following cLTD induction.

**Supplementary Table 2. Differentially expressed genes (DEGs) identified by TRAP-seq following cLTD.** Differential expression analysis of ribosome-associated transcripts comparing control and cLTD-treated neurons.

**Supplementary movie 1. Distinct mitochondrial fission dynamics during cLTP and cLTD.** Representative time-lapse recordings showing a rapid, transient burst of mitochondrial fission during cLTP and sustained, asynchronous fission during cLTD.

**Supplementary movie 2. Translation inhibition suppresses mitochondrial fission but not swelling during cLTD.** Representative time-lapse recordings comparing neurons acutely treated with DMSO or CHX. Note that translation inhibition markedly reduces mitochondrial fission, while transient mitochondrial swelling during cLTD persists.

**Supplementary movie 3. Local mitochondrial depolarization precedes fission during cLTD.** Representative time-lapse recording showing transient depolarization followed by fission of a single dendritic mitochondrion in a DIV7 immature hippocampal neuron

## Notes

### Competing Interest Statement

The authors have declared no competing interest.

### Summary of Updates

Addition of TRAP-seq to show an unbiased confirmation of upregulation of protein synthesis upon Long term Depression.

